# Bistability in Gene Regulation: Simulating Positive Feedback and Toggle Circuits Using Python and Hill Functions

**DOI:** 10.1101/2025.08.05.668746

**Authors:** Y K Yathu Krishna

## Abstract

Cellular decision-making relies heavily on bistable gene regulatory networks, which allow systems to respond to internal or external stimuli by switching between several stable expression states. Processes like cell differentiation, epigenetic memory, and the creation of artificial biological switches all depend on these dynamics. In this work, we introduce a simple and reproducible Python framework for modeling bistability in genetic feedback systems by means of ordinary differential equations (ODEs) driven by Hill functions. We employ two fundamental motifs, both of which are recognized for their ability to generate bistable behavior: a two-gene mutual inhibition toggle switch and a single-gene positive feedback loop. We investigate the effects of different Hill coefficients, production rates, and initial expression levels on system dynamics by numerical integration using SciPy. Our simulations show phase-plane convergence to several attractors, map expression outcomes over a grid of beginning circumstances, and illustrate the onset of bistability above a key Hill threshold. The delicate reliance of final states on cooperativity and beginning values is further demonstrated by heatmaps and bifurcation-like graphs. For accessibility, all code is hosted at Google Colab and is written in open-source Python. This study promotes research and teaching in synthetic biology, systems biology, and computational modeling while providing a simple yet effective computational framework for investigating the fundamentals of gene circuit bistability.

## 1. Introduction

A fundamental dynamic characteristic of biological systems is bistability, which allows cells to commit to discrete and stable states in the face of internal noise or variations in external inputs. Bipolarity, which is fueled by nonlinear themes like mutual inhibition and positive feedback, underpins processes like differentiation, toggle-like decisions, and memory in genetic regulatory networks [1]. To create resilient, programmable, and memory-retention-capable switch-like modules, modern systems and synthetic biology mostly rely on modeling these bistable structures [2,3]. Ordinary differential equations (ODEs) are frequently used to model gene circuits displaying bistability, and Hill-type kinetics is used to represent transcriptional regulation. In response to graded inputs, the Hill coefficient’s nonlinearity allows for switch-like transitions that resemble cooperative binding or extremely sensitive regulatory responses [4].

Understanding basic patterns like positive autoregulatory loops, in which a gene activates its own expression, and the genetic toggle switch, which consists of two mutually repressive genes, has been made possible thanks in large part to these models. Under certain parameter regimes, both architectures can produce bistable dynamics [5]. The use of bistable switches in artificial gene networks is still being expanded upon in recent research. Zhou et al. (2023), for instance, used stacked toggle switches to demonstrate programmable logic operations in mammalian cells, accomplishing multi-state control with bistable modules [6]. The translational importance of such circuits was also highlighted by Wang et al. (2021), who created an optogenetically controlled bistable toggle system to modify differentiation in stem cells [7]. The use of bistable switches in prokaryotic and eukaryotic systems has also been advanced by developments in synthetic transcriptional regulators and CRISPR-based repressors [8,9].

Computational simulation is still essential for examining the behavior of bistable systems prior to physical implementation, even with advancements in experimentation. High-level settings for biological circuit design are provided by programs like BioCRNpyler [10] and Tellurium [11], but there is still a demand for simple, lightweight, code-transparent frameworks that prioritize simplicity and reproducibility. Python is perfect for simulating the dynamics of genetic switches because of its easily navigable scientific stack (NumPy, SciPy, Matplotlib), particularly in educational and research contexts. In this work, we introduce a basic Python-based simulation framework for leveraging Hill-function ODEs to describe bistability in genetic regulatory systems.

We examine two canonical motifs: a two-gene toggle switch for mutual inhibition and a single-gene positive feedback loop. We model the time evolution of expression states using SciPy’s solve_ivp integrator, and assess the effects of initial conditions, production rate, and Hill coefficient on system behavior. Our findings illustrate hysteresis and switch-like transitions, show the evolution of bistability as a function of cooperativity, and use phase-plane trajectories and steady-state heatmaps to display attractor landscapes. Our work provides a pedagogically oriented and research-ready simulation framework that complements bigger modeling platforms by emphasizing transparency, modularity, and ease of use. To encourage open science and reproducibility, all code is written in Python and made accessible through an interactive Google Colab notebook. This work provides a rapid prototyping environment and a teaching tool for the study and creation of bistable gene circuits in synthetic biology.

## 2. Materials and Methods

### 2.1. Computational Environment

Python 3.10 on Google Colaboratory, a free cloud-based platform that facilitates repeatable, code-based computational experiments, was used for all simulations [12]. Because of its pre-installed scientific software for biological modeling, accessibility, and simplicity of sharing, Google Colab was selected [13]. The Python libraries listed below were utilized:

- NumPy (v1.24+) – for numerical computations and array handling [14]
- SciPy (v1.11+) – for solving ordinary differential equations using `solve_ivp` with Runge–Kutta integration [15]
- Matplotlib (v3.7+) – for plotting time series, phase-plane trajectories, and heatmaps [16]
- Seaborn (v0.12+) – for enhanced statistical visualization and heatmap rendering [17] All packages were either pre-installed in Google Colab or installed using `pip`.

### 2.2. Model Overview

To investigate how bistability arises in artificial gene networks, two basic gene regulatory motifs were modeled:

- A positive feedback loop is a self-reinforcing loop created when a gene product triggers its own transcription. [18]
- The Toggle transition is a two-gene mutual repression system that can transition between two states and store memory. [19]

Ordinary differential equations (ODEs) were used to implement these motifs, and Hill-type regulatory functions were used to mimic them dynamically.

### 2.3. Mathematical Formulation

i. Positive Feedback Model The autoregulatory feedback was described by:

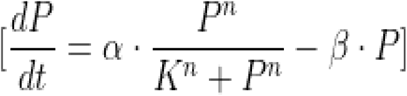 Where, *P*: Protein concentration α: Maximal production rate β: Degradation rate *K*: Activation threshold (half-maximal concentration) *n*: Hill coefficient (regulatory cooperativity) Such Hill-function-based positive feedback is a well-established approach to simulate ultrasensitive transcriptional regulation [20].
ii. Toggle Switch Model The mutual repression toggle switch was modeled as:

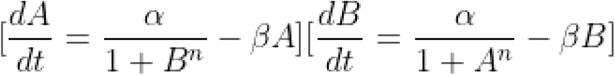 Where, *A,B* :Concentrations of the two repressors α: Maximal production rate β: Degradation rate *n*: Hill coefficient (regulatory cooperativity) This system exhibits bistability under appropriate cooperativity values and has been widely studied in synthetic biology [21,22].

### 2.4. Simulation Parameters

All simulations used the following baseline parameters unless otherwise specified:

| Parameter | Description | Value |
| --- | --- | --- |
| $\alpha$ | Max production rate | 5.0 |
| $\beta$ | Degradation rate | 1.0 |
| $K$ | Activation threshold (positive feedback only) | 1.0 |
| $n$ | Hill coefficient | 1 to 6 |
| $t$ | Time range | 0 to 50 units |
| $P0, A0, B0$ | Initial protein concentrations | Varied |

The adaptive Runge–Kutta (RK45) method, which is commonly used for biological simulations, was used to solve ODEs via scipy.integrate.solve_ivp [15,23]. The t_eval option was used to assess each simulation at 500 evenly spaced time periods.

### 2.5. Parameter Sweeps and Visualizations

The following simulation experiments were performed:

- Hill Coefficient Sweep: Simulated toggle switch behavior for *n* = 1 to 6, recording steady-state levels of *A and B* to observe the emergence of bistability [24].
- Initial Condition Grid Mapping: Varied *A*0 *and B*0 over a 25×25 grid to generate heatmaps showing regions of distinct final states (bistable attractor zones) [25].
- Phase-Plane Trajectories: Plotted *A vs B* over time from different initial conditions to visualize multistability and convergence [26].
- Bifurcation-Like Behavior: Final expression levels were plotted as a function of increasing *n*, revealing sharp transitions consistent with bifurcation thresholds [27].

Seaborn and matplotlib.pyplot were used to create all of the visualizations. heatmap with axis formatting and color schemes suitable for publication.

### 2.6. Code and Data Availability

Following preprint posting, all Python source code, simulation notebooks (.ipynb), and figure-generation scripts will be made publically accessible through GitHub. There will also be a link to Google Colab for real-time replication.

## 3. Results

### 3.1. Effect of Cooperativity on Positive Feedback Dynamics

To evaluate how cooperativity affects gene expression in a positive feedback loop, we simulated the time evolution of a single-gene autoregulatory system using varying Hill coefficients (*n*= 1, 2, 4,8). Increasing the Hill coefficient inhibits expression dynamics and ultimately results in ultrasensitive or switch-like behavior, as seen in **Figure 1**. The system shows a smooth ascent to a high steady-state level of protein expression at low cooperativity (*n*=1). Higher values of nnn, on the other hand, inhibit stable activation and cause a fast decay or suppression of expression, which is in line with the threshold effects seen in bistable systems.

**Figure 1.**
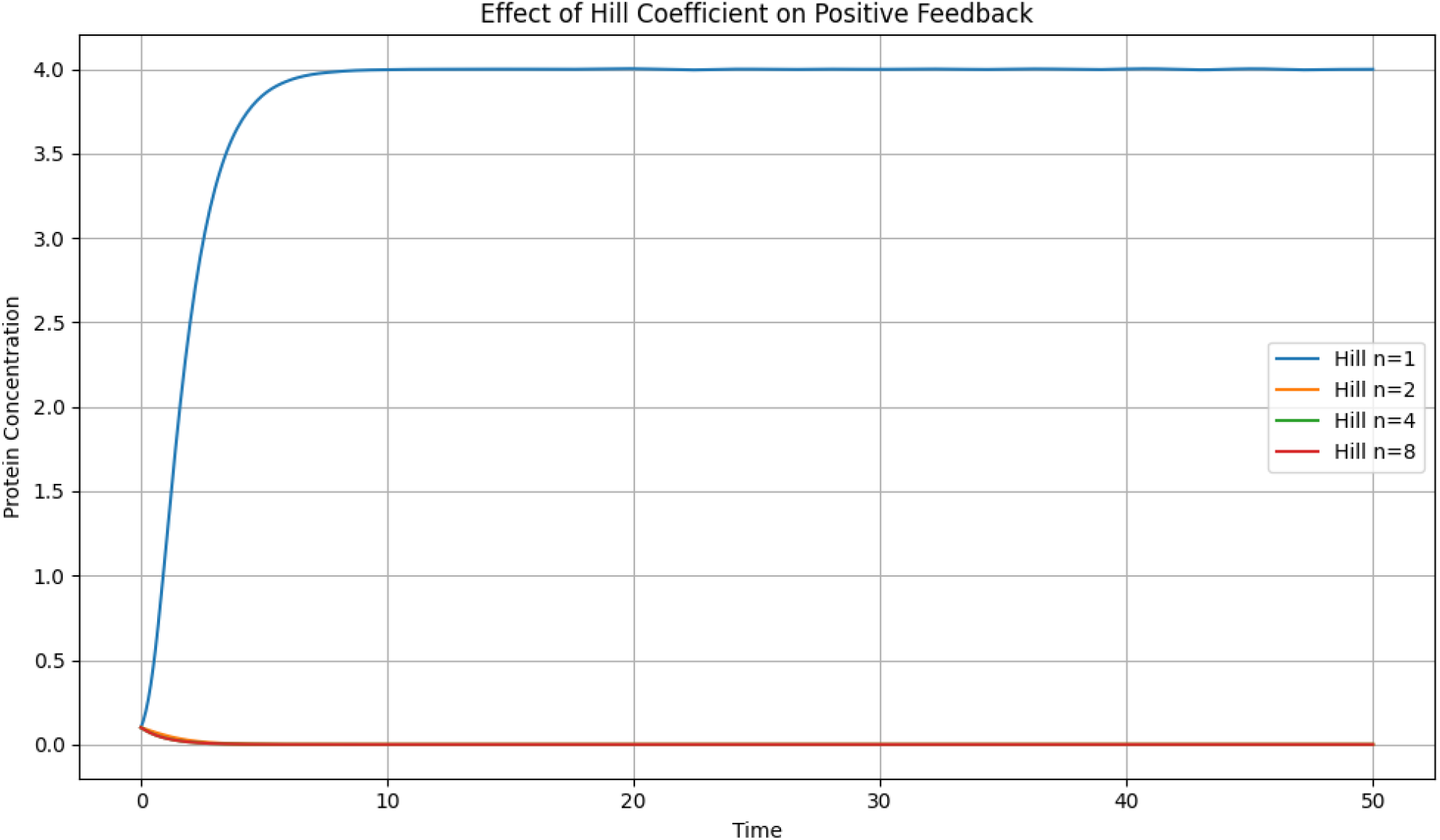

### 3.2. Dynamics of Mutual Repression in a Genetic Toggle Switch

The behavior of a canonical bistable motif, a two-gene mutual inhibition toggle switch, was then investigated. Time-course simulations with a fixed Hill coefficient of *n* = 4 were run from a variety of initial circumstances. One of two stable expression states is reached by the system, as illustrated in **Figure 2**: either gene A dominates while gene B is inhibited, or vice versa. The presence of numerous attractors is indicated by the outcome’s significant dependence on initial conditions. This is typical of synthetic gene circuits’ memory-storing and bistable switching behaviors.

**Figure 2.**
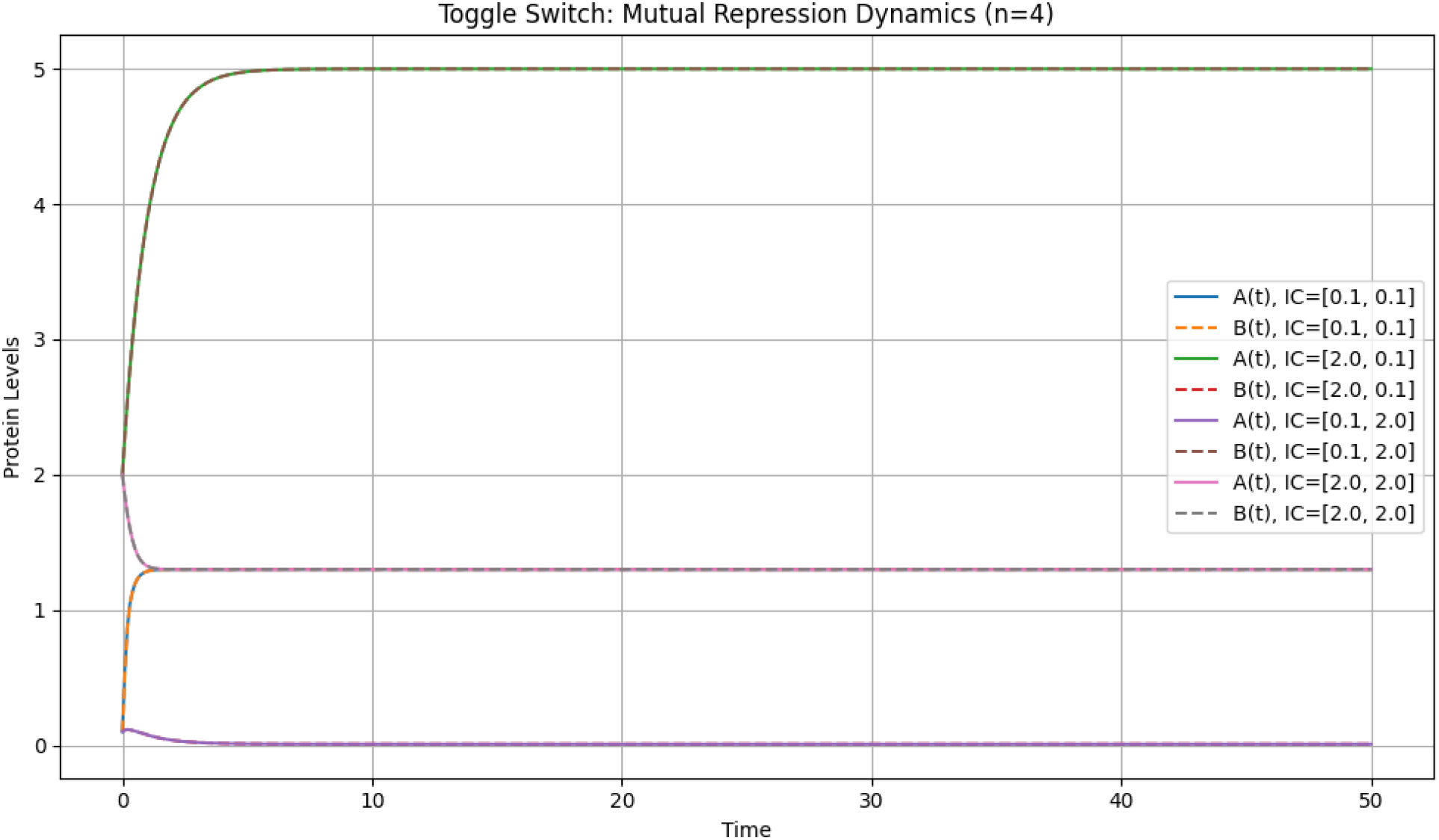

### 3.3. Phase-Plane Analysis Reveals Attractors and Trajectories

Phase-plane plots were created for the toggle switch model utilizing various initial condition combinations in order to show convergence toward stable expression levels. Trajectories in the A-B plane are shown in **Figure 3**, where they are clearly separated into separate attractor basins. While off-diagonal initial conditions quickly resolve into polarized expression levels of either A or B, solutions originating close to the diagonal exhibit delayed dynamics and convergence to a balanced state. The system’s bistable nature is further supported by the attractors’ symmetry.

**Figure 3.**
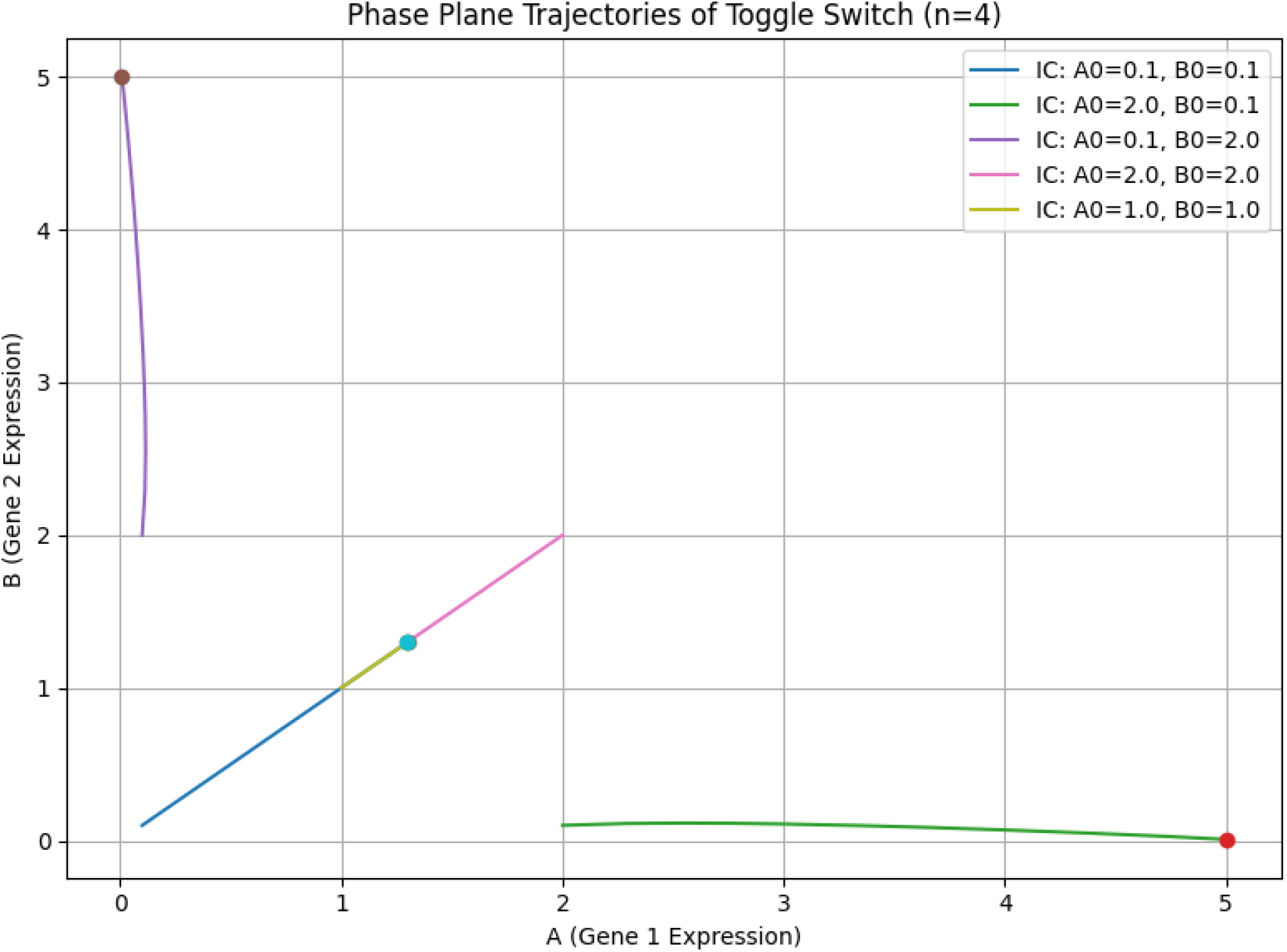

### 3.4. Emergence of Bistability with Increasing Hill Coefficient

We conducted a parameter sweep of the Hill coefficient *n* from 1 to 6 while keeping all other parameters fixed in order to evaluate how cooperativity drives bistability. The toggle switch system was set up with asymmetrical circumstances (*A*0 = 0.5, *B*0 = 2.0) for every value of *n*, and the expression levels at the end were noted. **Figure 4** illustrates how bistability becomes evident at *n* ≥2, with a distinct division between A and B’s final expression levels. The system stabilizes close to symmetric levels for low *n* indicating monostability. The bifurcation-like transition demonstrates that bistability requires coordinated repression.

**Figure 4.**
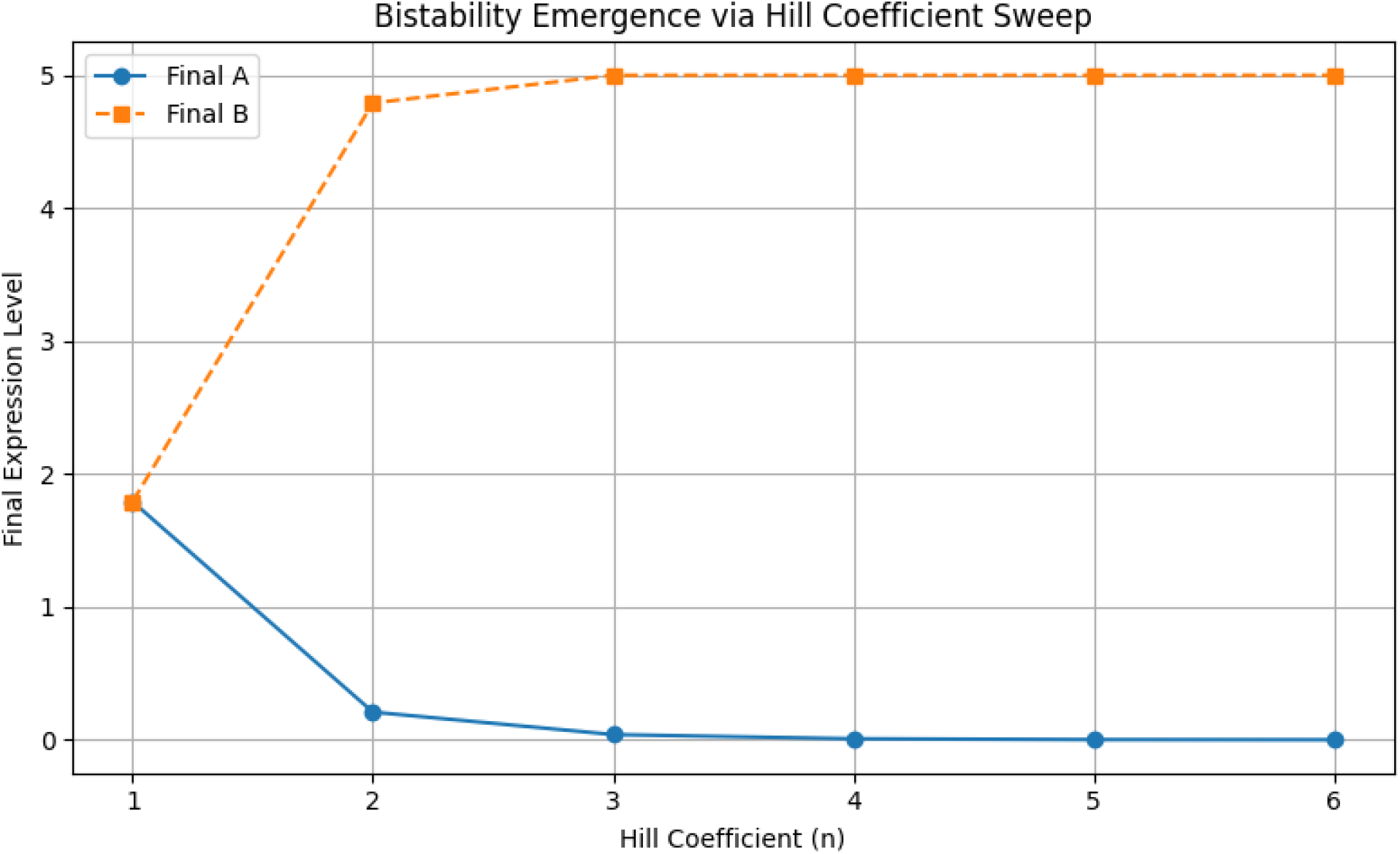

### 3.5. Bistability Landscape from Initial Condition Grid

We mapped the system’s steady-state result across a 25×25 grid of beginning conditions in order to further confirm bistability (*A*0, *B*0 ϵ [0. 1,3. 0]). **Figure 5** shows a heatmap depicting the final gene A concentration. A abrupt change from A-dominant to B-dominant expression states is shown by the diagonal boundary that divides the red and blue zones. This strongly implies bistable gene networks’ defining characteristics of multistability and sensitive dependency on beginning conditions.

**Figure 5.**
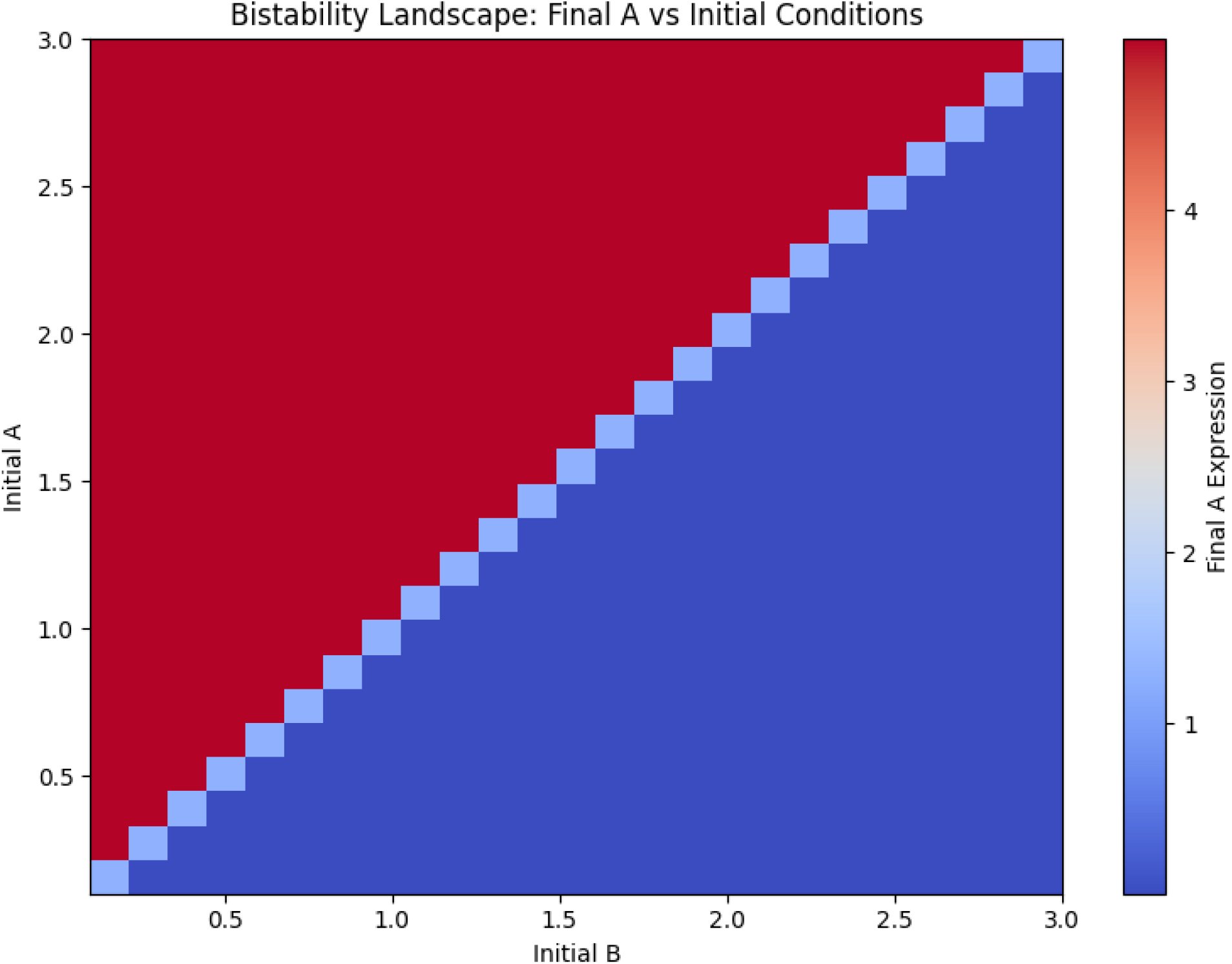

## 4. Discussion

In both natural and artificial gene regulatory networks, bistability is a basic emergent feature. Our simulations show that bistable expression profiles can arise under certain parameter regimes from minimum gene circuit topologies, specifically a mutual repression toggle switch and a single-gene positive feedback loop. We were able to investigate the threshold behavior that underlies multistability and model nonlinearity in regulatory dynamics by utilizing Hill functions with variable cooperativity. Our findings verify that the system’s responsiveness is sharpened by raising the Hill coefficient, allowing for sudden changes between low and high expression stages. This cooperative repression creates discrete attractor states in the toggle switch motif, the results of which depend on initial conditions. These results are in line with earlier research that emphasizes how ultrasensitivity drives switch-like dynamics [21,24]. The toggle switch’s bistable landscape is evident via phase-plane and heatmap visualizations, which show distinct borders between the A-dominant and B-dominant attractors. The competitive dynamics and underlying symmetry between mutually repressive regulators are reflected in this. The simulations also demonstrate hysteresis-like behavior, in which the system’s ultimate outcome is heavily influenced by its initial settings, emulating cellular memory in biological decision-making circuits. Crucially, Google Colab’s open-source Python tools were used for all simulations, guaranteeing accessibility and reproducibility for both researchers and students.

Our method provides a portable substitute for more complex simulation systems like Tellurium or COPASI [10,23], which makes it perfect for quick prototyping and teaching. A foundation for expanding minimum gene circuit models to incorporate delays, noise, stochastic effects, or spatial interactions is provided by this work. Similar ODE frameworks may be used in future research to examine CRISPR-based logic components, optogenetic toggles, or circuit design under metabolic limitations.

## 5. Conclusion

We introduce a computational framework based on Python that uses Hill-function ODEs to simulate bistable gene regulation systems. Our research shows how initial circumstances and cooperativity influence the dynamic behavior of genetic feedback circuits, allowing for multistability and toggle-like transitions. We verified the emergence of bistability in single-gene and two-gene systems using attractor heatmaps, phase-plane plots, Hill coefficient sweeps, and time-course simulations. This framework is ideal for teaching, outreach, and early synthetic circuit design because of its code’s readability and clarity. To ensure maximum reproducibility, all simulations were executed on Google Colab with openly available software. We anticipate that our study will help develop a valuable research and teaching tool for comprehending genetic switches and creating reliable artificial biological networks.

## Supporting information

Supplementary Codes

## Notes

https://github.com/ykayuwu/Bistability-Project-Code

### Competing Interest Statement

The authors have declared no competing interest.

