## Supplementary Codes for "Bistability in Gene Regulation: Simulating Positive Feedback and Toggle Circuits Using Python and Hill Functions"

```
import numpy as np
from scipy.integrate import solve_ivp
import matplotlib.pyplot as plt

# Model 1: Positive Autoregulation (Self-activation)
def positive_feedback(t, P, alpha=5.0, K=1.0, n=2, beta=1.0):
    # dP/dt = α * (P^n / (K^n + P^n)) - β * P
    dPdt = alpha * (P[0]**n / (K**n + P[0]**n)) - beta * P[0]
    return [dPdt]

# Time range
t_span = (0, 50)
t_eval = np.linspace(*t_span, 500)

# Initial concentration
P0 = [0.1]

# Try different Hill coefficients
hill_coeffs = [1, 2, 4, 8]

plt.figure(figsize=(10, 6))

for n in hill_coeffs:
    sol = solve_ivp(positive_feedback, t_span, P0, args=(5.0, 1.0, n, 1.0), t_eval=t_eval)
    plt.plot(sol.t, sol.y[0], label=f'Hill n={n}')

plt.title('Effect of Hill Coefficient on Positive Feedback')
plt.xlabel('Time')
plt.ylabel('Protein Concentration')
plt.legend()
plt.grid(True)
plt.tight_layout()
plt.show()
```

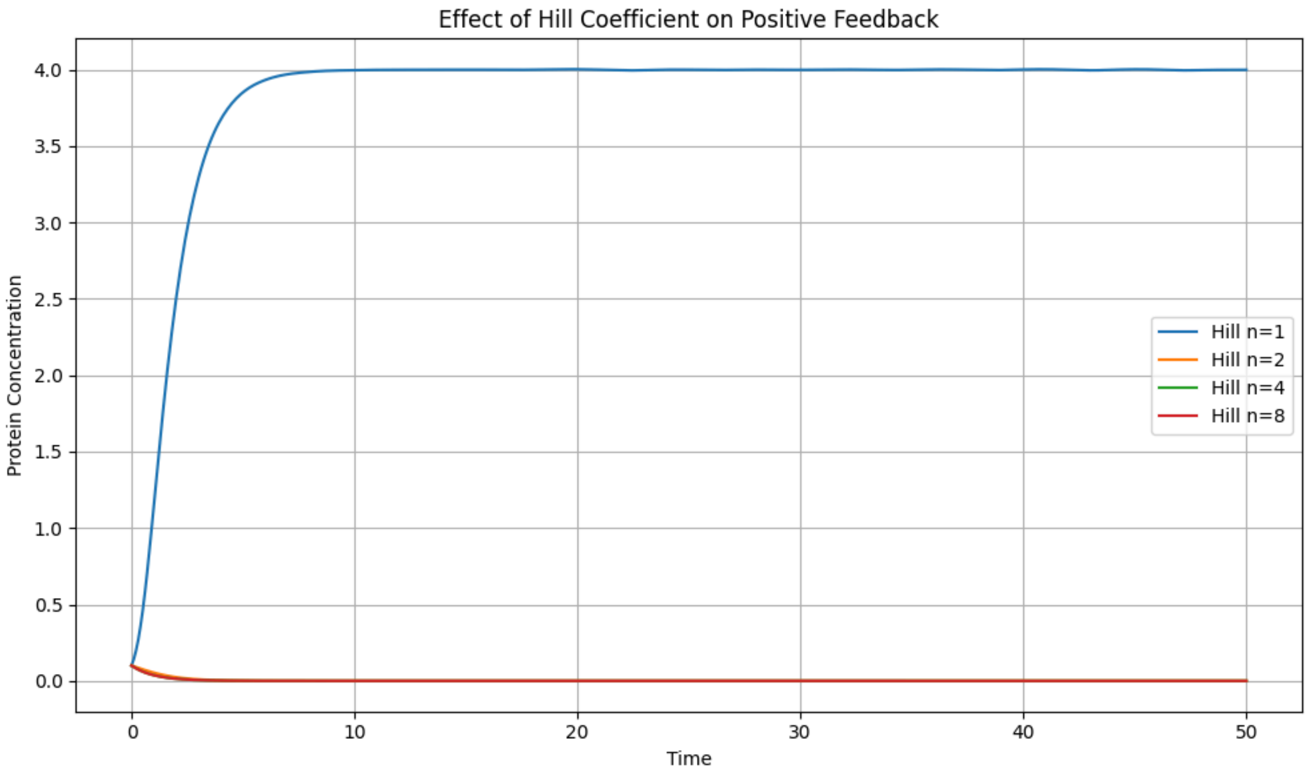


```
import numpy as np
from scipy.integrate import solve_ivp
import matplotlib.pyplot as plt

# Toggle Switch Model: Two genes mutually repress each other
def toggle_switch(t, y, alpha=5.0, beta=1.0, n=2):
    A, B = y
    dA_dt = alpha / (1 + B**n) - beta * A
    dB_dt = alpha / (1 + A**n) - beta * B
    return [dA_dt, dB_dt]

# Simulation time
t_span = (0, 50)
```

```
t_eval = np.linspace(*t_span, 500)

# Try different initial conditions to explore bistability
initial_conditions = [
    [0.1, 0.1],
    [2.0, 0.1],
    [0.1, 2.0],
    [2.0, 2.0]
]

# Run simulation and plot results
plt.figure(figsize=(10, 6))

for ic in initial_conditions:
    sol = solve_ivp(toggle_switch, t_span, ic, args=(5.0, 1.0, 4), t_eval=t_eval)
    plt.plot(sol.t, sol.y[0], label=f"A(t), IC={ic}")
    plt.plot(sol.t, sol.y[1], '--', label=f"B(t), IC={ic}")

plt.title('Toggle Switch: Mutual Repression Dynamics (n=4)')
plt.xlabel('Time')
plt.ylabel('Protein Levels')
plt.legend()
plt.grid(True)
plt.tight_layout()
plt.show()
```

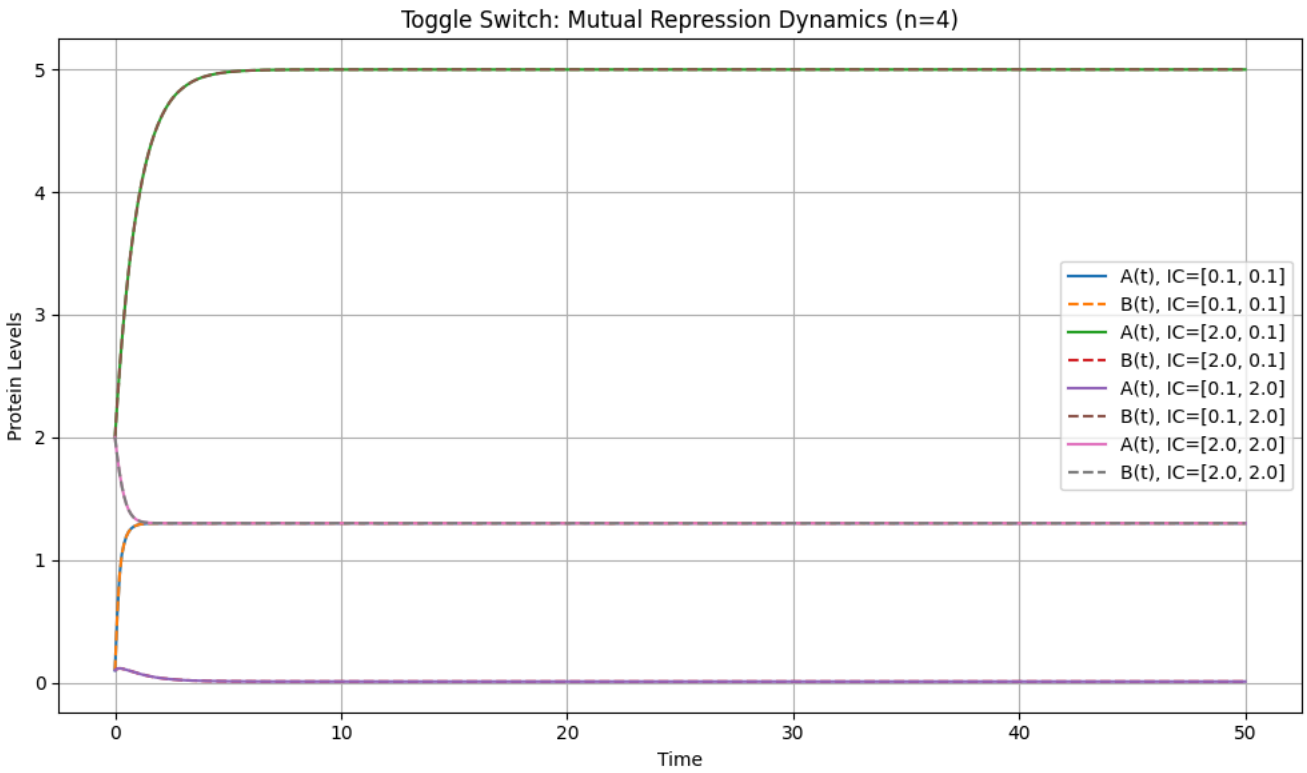


```
import numpy as np
from scipy.integrate import solve_ivp
import matplotlib.pyplot as plt

# Toggle Switch ODEs
def toggle_switch(t, y, alpha=5.0, beta=1.0, n=4):
    A, B = y
    dA_dt = alpha / (1 + B**n) - beta * A
    dB_dt = alpha / (1 + A**n) - beta * B
    return [dA_dt, dB_dt]

# Time range
t_span = (0, 50)
t_eval = np.linspace(*t_span, 500)

# Initial conditions
initial_conditions = [
    [0.1, 0.1],
    [2.0, 0.1],
    [0.1, 2.0],
    [2.0, 2.0],
    [1.0, 1.0]
]
```

```
# Phase-plane plot
plt.figure(figsize=(8, 6))

for ic in initial_conditions:
    sol = solve_ivp(toggle_switch, t_span, ic, args=(5.0, 1.0, 4), t_eval=t_eval)
    A, B = sol.y
    plt.plot(A, B, label=f'IC: A0={ic[0]}, B0={ic[1]}')
    plt.plot(A[-1], B[-1], 'o')  # Mark final point

plt.title('Phase Plane Trajectories of Toggle Switch (n=4)')
plt.xlabel('A (Gene 1 Expression)')
plt.ylabel('B (Gene 2 Expression)')
plt.grid(True)
plt.legend()
plt.tight_layout()
plt.show()
```

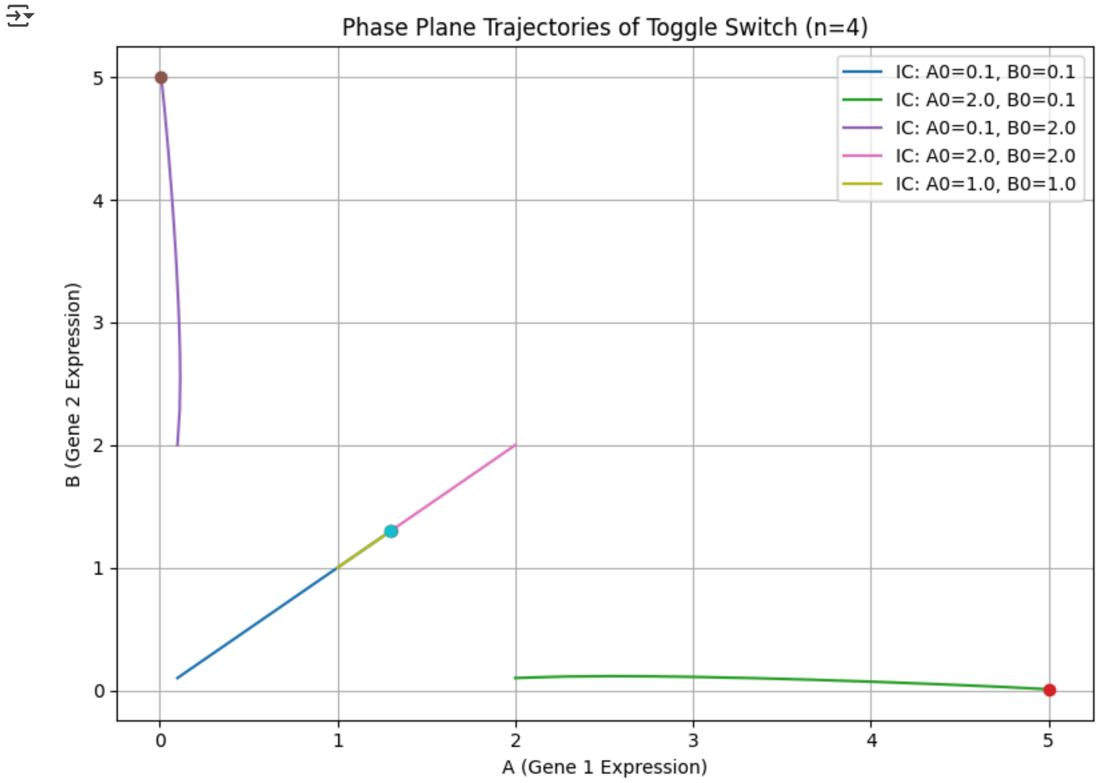


```
import numpy as np
from scipy.integrate import solve_ivp
import matplotlib.pyplot as plt

# Toggle switch system
def toggle_switch(t, y, alpha=5.0, beta=1.0, n=2):
    A, B = y
    dA_dt = alpha / (1 + B**n) - beta * A
    dB_dt = alpha / (1 + A**n) - beta * B
    return [dA_dt, dB_dt]

# Time span and evaluation
t_span = (0, 50)
t_eval = np.linspace(*t_span, 500)

# Initial condition
IC = [0.5, 2.0]  # Start slightly asymmetric

# Hill coefficients to try
n_values = np.arange(1, 7)
A_final = []
B_final = []

for n in n_values:
    sol = solve_ivp(toggle_switch, t_span, IC, args=(5.0, 1.0, n), t_eval=t_eval)
    A_final.append(sol.y[0][-1])
    B_final.append(sol.y[1][-1])

# Plot final steady-state levels vs Hill coefficient
plt.figure(figsize=(8, 5))
plt.plot(n_values, A_final, 'o-', label='Final A')
```

```
plt.plot(n_values, B_final, 's--', label='Final B')
plt.title('Bistability Emergence via Hill Coefficient Sweep')
plt.xlabel('Hill Coefficient (n)')
plt.ylabel('Final Expression Level')
plt.grid(True)
plt.legend()
plt.tight_layout()
plt.show()
```

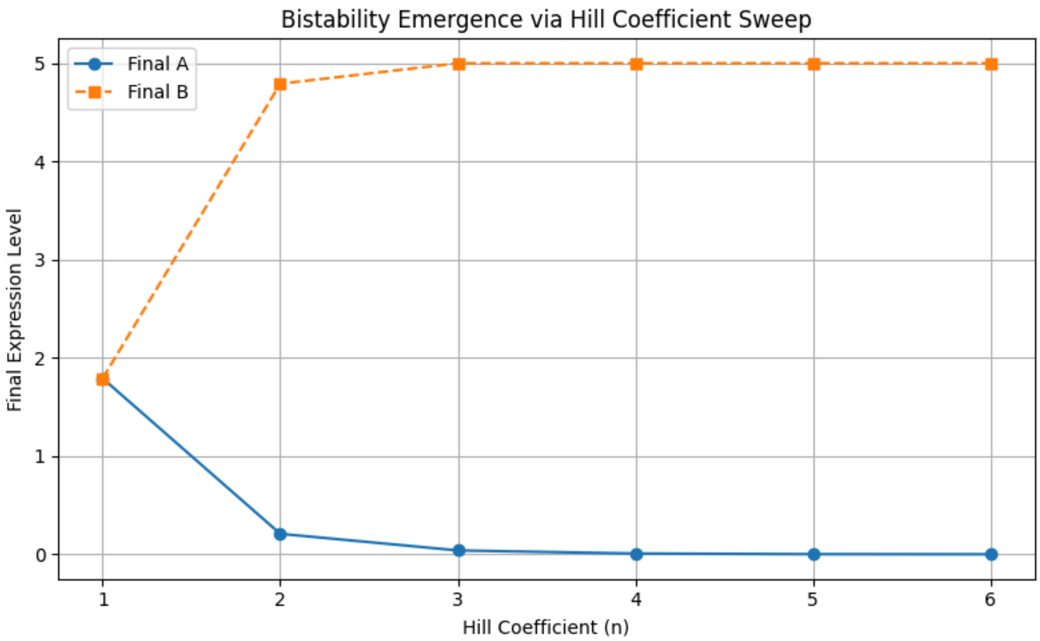


```
import numpy as np
import matplotlib.pyplot as plt
from scipy.integrate import solve_ivp

# Toggle switch ODE
def toggle_switch(t, y, alpha=5.0, beta=1.0, n=4):
    A, B = y
    dA_dt = alpha / (1 + B**n) - beta * A
    dB_dt = alpha / (1 + A**n) - beta * B
    return [dA_dt, dB_dt]

# Time and evaluation settings
t_span = (0, 50)
t_eval = np.linspace(*t_span, 300)

# Grid of initial A and B values
a0_vals = np.linspace(0.1, 3.0, 25)
b0_vals = np.linspace(0.1, 3.0, 25)
A_matrix = np.zeros((len(a0_vals), len(b0_vals)))

# Run simulations
for i, a0 in enumerate(a0_vals):
    for j, b0 in enumerate(b0_vals):
        sol = solve_ivp(toggle_switch, t_span, [a0, b0], args=(5.0, 1.0, 4), t_eval=t_eval)
        A_matrix[i, j] = sol.y[0][-1]  # final A value

# Plot heatmap
plt.figure(figsize=(8, 6))
plt.imshow(A_matrix, extent=[b0_vals[0], b0_vals[-1], a0_vals[0], a0_vals[-1]],
           origin='lower', aspect='auto', cmap='coolwarm')
plt.colorbar(label='Final A Expression')
plt.title('Bistability Landscape: Final A vs Initial Conditions')
plt.xlabel('Initial B')
plt.ylabel('Initial A')
plt.tight_layout()
plt.show()
```

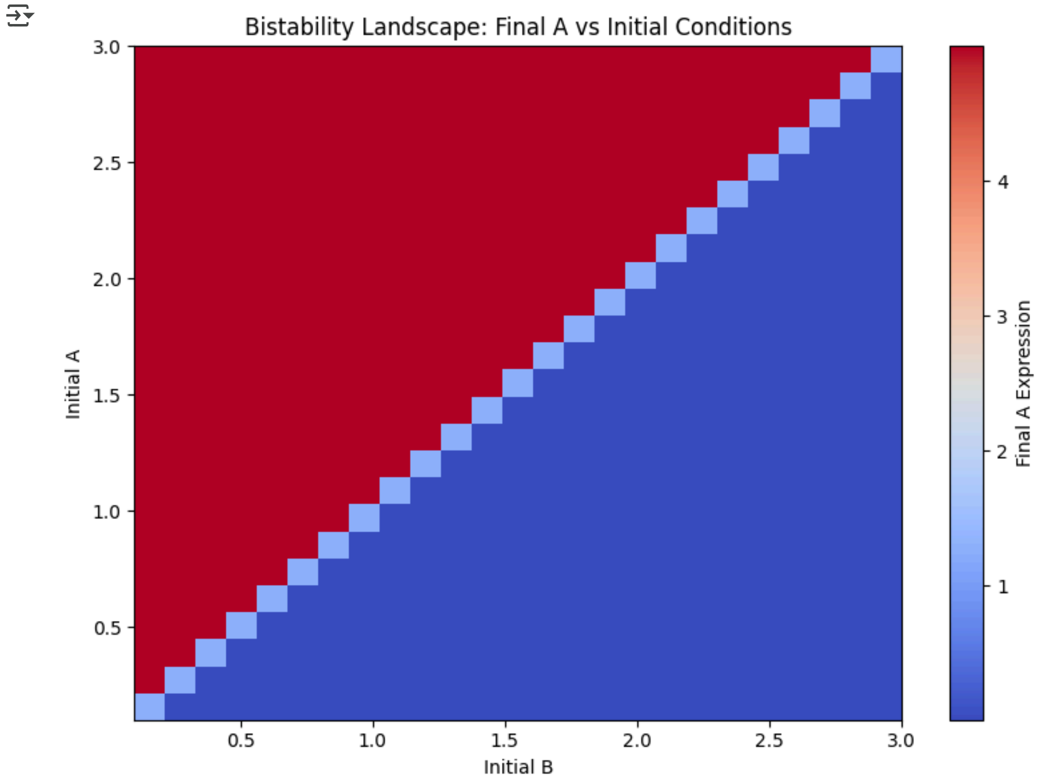

Bistability Landscape: Final A vs Initial Conditions
Initial A
Initial B
Final A Expression
3.0
2.5
2.0
1.5
1.0
0.5
4
3
2
1
